# Media representation of spiders may exacerbate arachnophobic sentiments by framing a distorted perception of risk

**DOI:** 10.1101/2020.04.28.065607

**Authors:** Stefano Mammola, Veronica Nanni, Paolo Pantini, Marco Isaia

**Affiliations:** Molecular Ecology Group (MEG), Water Research Institute, National Research Council of Italy (CNR-IRSA), Largo Tonolli 50, 28922 Verbania Pallanza, Italy; Department of Earth, Environmental and Life Science (DISTAV), University of Genoa, Via Balbi, 5, 16126 Genova, Italy; Museo civico di Scienze Naturali “E. Caffi.”, Piazza Cittadella 10, 24129, Bergamo, Italy; Department of Life Sciences and Systems Biology, University of Turin, Via Accademia Albertina 13, 10123 Torino, Italy

**Keywords:** Arachnophobia, Black widows, Emotional contagion, Envenomation, Facebook, Fake news, Latrodectism, Loxoscelism, Mass media, Recluse spiders, Social media, Spider bite

## Abstract

Spiders are able to arouse strong emotional reactions in humans. While spider bites are statistically rare events, our perception is skewed towards the potential harm spiders can cause to humans. We examined the human dimension of spiders through the lens of traditional media, by analysing more than 300 spider-related news published online in Italian newspapers between 2010 and 2020. We observed a recent, exponential increase in the frequency of the news, particularly those focused on medically important spiders – the Mediterranean black widow (*Latrodectus tredecimguttatus*) and the Mediterranean recluse (*Loxosceles refescens*). The news quality was generally poor: 70% contained different types of error, 32% were exaggerated, and in virtually none was an expert consulted. Overstated news referring to spider bites were significantly more shared on social media, thus contributing to frame a distorted perception of the risk associated with a spider bite and possibly reducing general public tolerance of spiders.

## INTRODUCTION

Wildlife is an important emotional trigger in humans (Jacobs, 2009, 2012; Hicks & Stewart, 2018). Admiration and respect, surprise and excitement, transcendent feelings, but also fear and disgust are just a few examples illustrating the spectrum of emotions reported by people experiencing encounters with wildlife (Hicks & Stewart, 2018). An interesting aspect of the human dimension of wildlife is that sensitivity toward animals is largely conserved in most contemporary societies, even though wildlife no longer plays a central role in our every-day lives (Franklin & White, 2001). Studies suggest that emotional feelings toward wildlife are, indeed, in-born (Strommen, 1995; Davey et al., 1998; Prokop & Tunnicliffe, 2008; DeLoache, Pickard, & LoBue, 2010), often recurring with striking similarities across diverse cultural settings (Davey et al., 1998). As a direct consequence, animals-related emotions end up playing a key role in scientific and socio-political debates around both the management and conservation of wildlife (Jones, 2006; Singh, 2009; Frank, Johansson, & Flykt, 2015; Zainal Abidin & Jacobs, 2019; Drijfhout, Kendal, & Green, 2020; Straka, Miller, & Jacobs, 2020), and in the perception of risk (Knopff, Knopff, & St. Clair, 2016; Hathaway et al., 2017; Bombieri et al., 2018; Nanni et al., 2020).

Spiders are iconic examples of animals that can bring about strong emotional reactions in humans (Michalski & Michalski, 2010; Lemelin & Yen, 2015; Hauke & Herzig, 2017; Mammola, Michalik, Hebets, & Isaia, 2017), leading to a distorted perception of risk, especially when referring to spider bites. While less than 0.5% of spider species are capable of causing severe envenomation in humans (Hauke & Herzig, 2017), and no proven fatality due to spider bites had occurred in the past few decades (Nentwig & Kuhn-Nentwig, 2013; Nentwig, Gnädinger, Fuchs, & Ceschi, 2013; Stuber & Nentwig, 2016), the perception of the risk associated with spider bites remains skewed towards the potential harm spiders can cause in humans (Hauke & Herzig, 2017). These feelings seemingly find their psychological roots in our ancestral fear of venomous animals (Knight, 2008; Gerdes, Uhl, & Alpers, 2009), but might also have a cultural component (Davey, 1994; Merckelbach, Muris, & Schouten, 1996; Davey et al., 1998). As Cavell (2018, p. 2) nicely put it “… *one of the most remarkable aspects of modern human-spider relations is the prevalence of arachnophobia in places with few or no highly dangerous spider species*”. Indeed, even though human-spiders encounters are frequent events because spiders are omnipresent in all terrestrial ecosystems (Turnbull, 1973), including indoor environments (Bertone et al., 2016), the objective risk of being bitten by a harmful spider is minimal in most areas of the world (Diaz & Leblanc, 2007). These considerations raise the questions of why such a skewed perception of risk persists in modern societies (Lemelin & Yen, 2015).

It is known that humans have the tendency to evaluate risk through feelings and emotions rather than objectively (Slovic & Peters, 2006), often overestimating the frequency of statistically rare events. For example, many people fear flying, even though the casualties associated with civil flights are estimated to be in the order of 0.07 deaths per billion passenger miles (Savage, 2013). The same line of reasoning can be applied to people’s risk judgments of low probability events related to wildlife, such as being attacked by a large carnivore (Bombieri et al., 2018) or stung or bitten by a venomous animal (Langley, 2005).

Furthermore, a distorted perception of risk can be exacerbated by the way in which information is framed in the scientific literature (Bennett & Vetter, 2004; Stuber & Nentwig, 2016) or in traditional media sources (Gerber, Burton-Jeangros, & Dubied, 2011). As far as spiders are concerned, it has been demonstrated that there is a significant overdiagnosis of spider bites and envenomation in the medical literature (White, 2003; Bennett & Vetter, 2004; Vetter, 2004; Vetter et al., 2005; Vetter, Hinkle, & Ames, 2009; Stuber & Nentwig, 2016). A recent major role in spreading falsehoods about spiders could also be associated with traditional and social media, due to their high efficiency in conveying a message more directly and reaching a wider audience (Vosoughi, Roy, & Aral, 2018). It is understood how the media play an important role in the construction and circulation of risk images associated with animals, contributing to develop fears and ambivalence (Gerber et al., 2011). Yet, while spiders are the quintessential feared animals, there is still poor understanding of the role of the media in spreading (mis)information about them (Cushing & Markwell, 2010).

Here, we explored the human dimension of spiders in Italy through the lens of traditional and social media. We examined the media representations of human-spider encounters as published in Italian online newspapers over the past 10 years, in order to assess the accuracy, spreading, and sensationalistic content of news. We tackled the following questions:

i. What is the content and quality of the information of each spider-related media report?
ii. What is the temporal distribution of spider-related news?
iii. Which factors determine the effective spreading of news on social media?

Our over-arching goal is to understand the potential role of online media in exacerbating arachnophobic sentiments and promoting a distorted perception of the risk associated with spider bites. This is important given that these negative sentiments may ultimately lead to lowering public tolerance towards spiders and reducing conservation efforts towards them (Knight, 2008; Simaika & Samways, 2018).

## METHODS

### Media report search

We adapted the methodology of Bombieri et al. (2018) for retrieving media reports on human-spider encounters published in Italian online newspapers (Figure 1a). We carried out online searches in Italian with *Google news*, choosing multiple keyword combinations. We first searched for the Italian words for bite (“morso”), followed by spider (“ragno”) and one of the years between 2010 and 2020 (e.g., “morso ragno 2014”). We repeated the search using the word sting (“puntura”) instead of bite, given that it is frequently used (incorrectly) by journalists (among others; see, e.g., Afshari, 2016). We then repeated the search, changing the noun “ragno” (spider) to the Latin and vernacular names of spider species generally perceived as dangerous in Italy: *Cheiracantium punctorium* (“Ragno dal sacco giallo”), *Latrodectus tredecimguttatus* (“Argia”, “Malmignatta”, “Vedova nera”), *Loxosceles rufescens* (*“*Reclusa”, “Ragno eremita”, “Ragno violino”), and *Zoropsis spinimana* (“Falsa licosa”). We compiled the list of species based on our experience in years of interaction with the staff of the Anti-poison Center in Milan (Centro Antiveleni) and the San Giovanni Molinette hospital in Turin, who regularly contacted us asking for expert opinions on spider identification (on average 4.6 requests/month in 2019).

**Figure 1.**
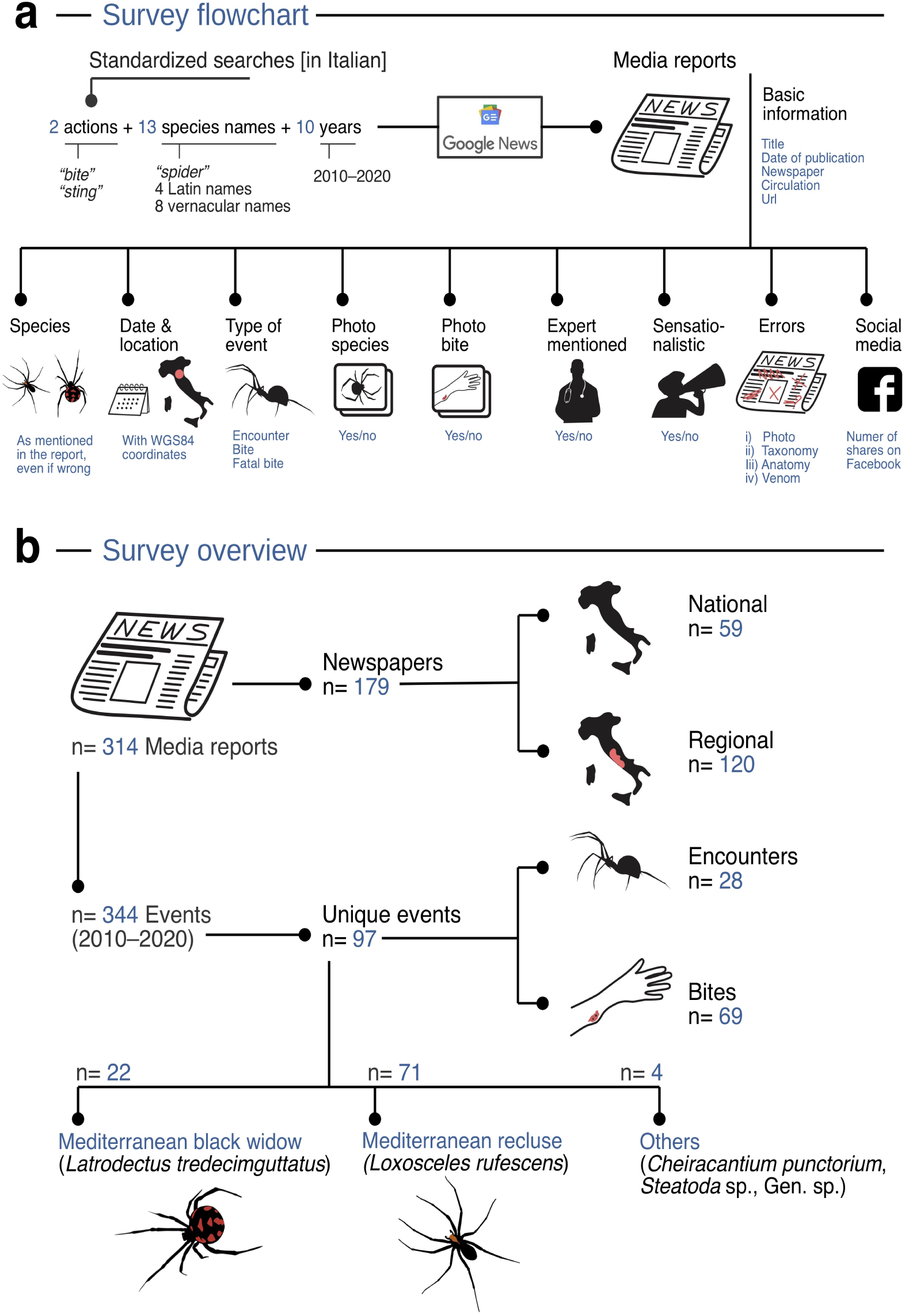
Infographic illustrating the study design and summary statistics: **a**) flowchart of the general methodology for retrieving media reports and mining relevant information; **b**) survey summary statistics.

This search strategy led to a total of 260 searches: 2 actions (“morso” or “puntura”) x 13 species names (the general words “ragno”, 4 Latin, and 8 vernacular species names) x 10 years (2010–2020). For each unique keyword search, we checked news up to the final available page in *Google news*, collecting all the media reports referring to one or more encounters in Italy between humans and spiders. We disregarded: i) media reports which did not mention a specific locality for the event; ii) media reports referring to spider bite events that occurred outside Italy (e.g., a report written in Italian but focusing on a spider bite that occurred in England); and iii) media reports not specifically reporting a spider-human encounter (e.g., news discussing best practices to deal with a spider bite).

### Media report content

For each media report, we first extracted basic information: a) title, b) date of publication, c) journal name, and d) journal circulation (‘Regional’ or ‘National”). We classified newspapers circulation as ‘Regional’ if their total circulation was below 50,000 copies and as ‘National’ if it was above 50,000 copies, using the 2017 Assessment for Press Circulation provided by the society Accertamenti Diffusione Stampa (ADS) srl. Whenever newspapers were not covered in this report, we used the information found on each newspaper’ webpage.

Then we read the full article and scored the e) spider species as it was mentioned in the media report (even if the species attribution was incorrect based on indirect evidence), f) type of event (“encounter”, “bite”, or “deadly bite”), g) year of the event, h) location of the event, i) presence/absence of photographs of the spider, j) presence/absence of photographs of the bite, and k) possible mention of an expert-opinion (doctor, arachnologist, or general biologist). Since several media reports were discussing the same event, we created an identifier for each unique event (“Event_ID”), by combining location and year of the event (e.g., “Terni_2018”). We also derived WGS84 coordinates for each event location, by geo-referencing the nearest city on *Google Earth*.

Following Nanni et al. (2020), we expressed the success of each media report as its spreading on social media, using the number of total shares in Facebook. We chose Facebook, as it is one of the most used social media platforms in Social Science research (e.g., Wilson, Gosling, & Graham, 2012; Kramer, Guillory, & Hancock, 2014). We extracted Facebook shares using the API tool available on ShareCount webpage (www.sharedcount.com; accessed on 2 March 2020). When the number of shares exceeds 999, this tools returns a rounded number (e.g., 1K for number of shares between 1000 and 1999). In such cases, we used the lowest number (1000). Even though we compared the number of shares for media reports published in different years, we consider this a reliable approach (see Nanni et al., 2020). Indeed, the share of online news on social media typically reaches a stable plateau at 30 days after publication (Papworth et al., 2015).

### Scientific quality of the media reports

We assessed the quality of each media report by checking for the presence/absence of four types of errors in text and figures:

i. errors in photographs, when the photograph(s) of the species in the media report (if any) did not correspond to the species mentioned in the text, or when the attribution was not possible (e.g., blurry photographs);
ii. errors in systematics and taxonomy, like the common mistake of considering spiders “insects” (Jambrina, Vacas, & Sánchez-Barbudo, 2010), but also subtle inaccuracies in term of Linnaean taxonomic ranks [e.g., Report_ID 271 (translated): “*…* the *‘malmignatta*’, a *genus* of Italian spider belonging to the *family* of the *species* of the black widow”];
iii. errors in venom and other physiological or medical aspects or terminology [e.g., Report_ID 147 (translated): “… the *venom sac* was removed with surgery”]; and
iv. errors in morphology and anatomy, such as the frequent “spider sting” instead of “spider bite” (Afshari, 2016).

Each error type was scored as present or absent, thus we did not counted cumulative errors of the same type in the same report.

### Classification of Sensationalism

Three authors (MI, SM, and VN) independently evaluated the title, subheadings, and main text of each media report, and assessed it as overstated (sensationalistic) or not (neutral). We took the consensus between the three independent evaluations to minimize the effect of subjectivity. Sensationalism in animal-related media reports is often associated with emotional words and expressions (Bombieri et al., 2018; Nanni et al., 2020). In our case, frequent words associated with sensationalistic content were alarm (“allarme”), agony (“agonia”), attack (“attacco”), devil (“diavolo”), fear (“paura”), hell (“inferno”), killer (“assassino”), nightmare (“incubo”), panic (“panico”), terrible (“terribile”), and terror (“terrore”). Examples of titles (literally translated from Italian) of sensationalistic versus non sensationalistic media reports focusing on the same Event_ID are, respectively: i) “[…] Sardinia and the nightmare of venomous spiders” versus “Black widow spider spotted in Sardinia, but the expert is happy: it is an indicator of biodiversity”; ii) “Alarm in Rome: Violin spiders strike again and again. Boom of hospitalisations” versus “Bitten by a violin spider, he was immediately hospitalized”; or iii) “Attacked by a violin spider, traffic warden miraculously survived” versus “Be aware of the violin spider: if it bites you, it can be dangerous”.

### Data analysis

We conducted all analyses in R (R Core Team, 2018). We graphically explored the content of media reports with barcharts and boxplots with ‘ggplot2’ (Wickham, 2016). For the two most abundant species, *Latrodectus tredecimguttatus* and *Loxosceles rufescens*, we explored temporal distribution of media reports using density plot, by computing a kernel density estimate with a 1.5 bandwidth adjustment for both the annual and monthly distribution of media reports (Wickham, 2016). For this and the following analysis, we excluded media reports published in 2020 given this year was covered only up to February.

We used generalized linear mixed models (GLMM) to explore the factors driving the share of news on Facebook. We followed Zuur & Ieno’s (2016) protocol for presenting regression-type analyses, whereby we: i) conducted data exploration and identified the dependency structure in the data; ii) explained, fitted, and validated the regression models; and iii) interpreted the regression output and presented the main effect plots.

The data exploration revealed the presence of four outliers in the number of shares, namely media reports shared over 15,000 times on Facebook. We removed these four observation from the database. Furthermore, we observed that 39.5% of media reports were never shared on Facebook (Figure 2b). However, since these are “true zeros” (*sensu* Blasco-Moreno, Pérez-Casany, Puig, Morante, & Castells, 2019), we did not apply zero-inflated models.

**Figure 2.**
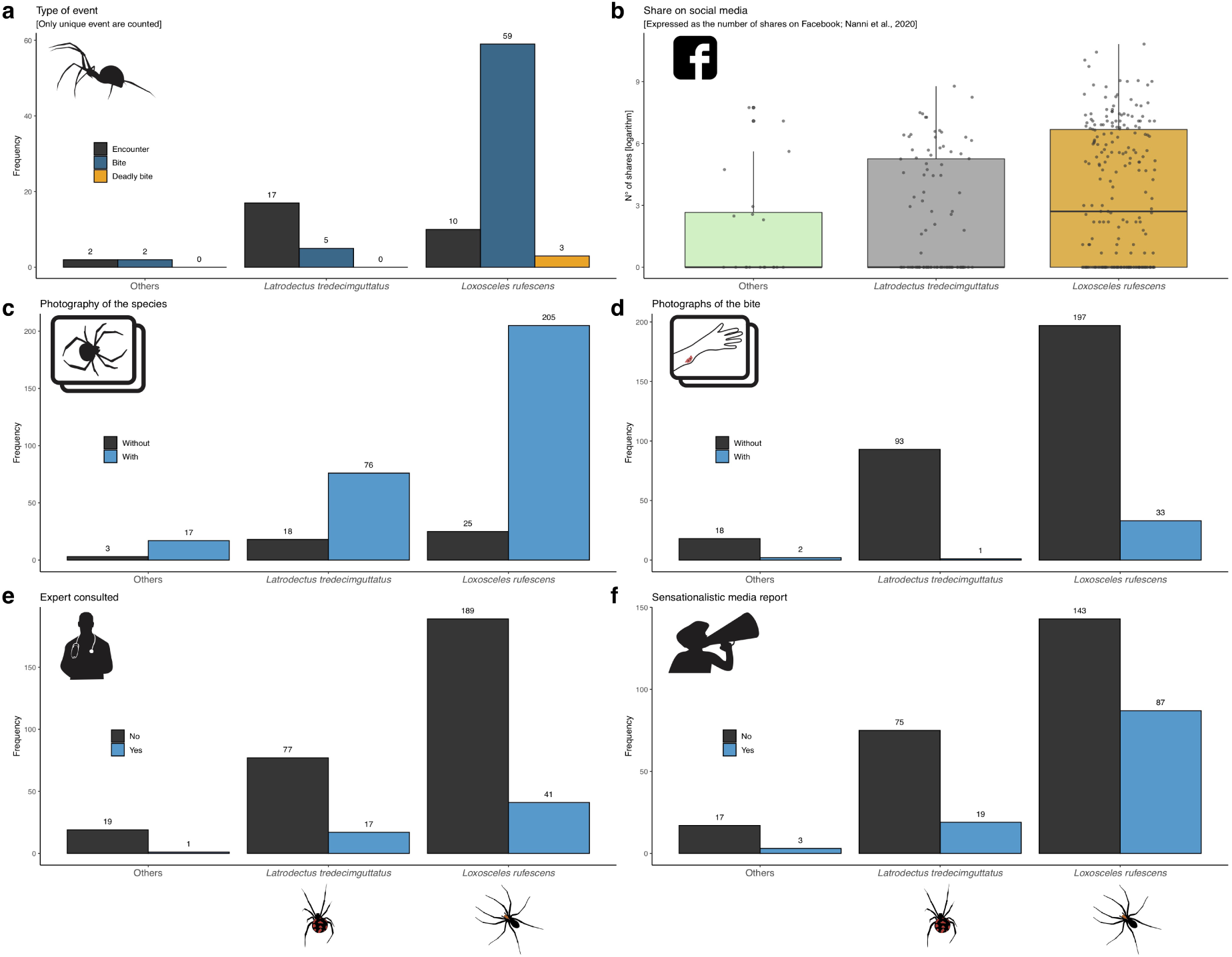
Content of media reports: **a**) Type of event covered by media reports focusing on *Latrodectus tredecimguttatus, Loxosceles rufescens*, and other species. **b**) Logarithm of total number of shares on Facebook (the grey dots are jittered observed values, whereas the boxplots summarize median, quantiles, and range). **c**) Frequency of species photographs in media reports. **d**) Frequency of bite photographs in media reports. **e**) Frequency of expert consultancy in media reports. **f**) Frequency of sensationalistic versus non-sensationalistic media reports.

We fitted GLMMs with ‘glmmADMB’ (Fournier et al., 2012), starting from an initial structure that included all covariates and random terms of interest:

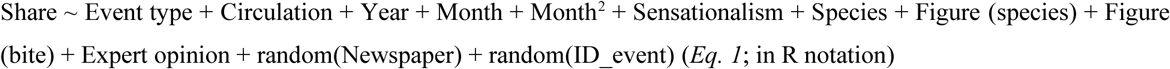

The random factor ‘Newspaper’ was introduced because reports published in the same newspaper usually share a similar language, style, and graphical elements. The random factor ‘Event_ID’ was introduced to take into account the fact that multiple reports in our dataset discussed the same events. We included the square of month (term month^2^) to capture a possible seasonal response of the shares during the year (i.e., a quadratic relationship between shares and month).

The numbers of Facebook shares are counts, so we initially chose a Poisson distribution. The Poisson GLMM was, however, highly over-dispersed (χ^2^: 227751553743; p<0.001) and so we switched to a negative binomial distribution. Once the initial model had been fitted, we performed a step-wise model selection in ‘MuMIn’ (Bartoń, 2019). We based the model reduction on Aikaike Information Criterion (AIC) and Aikaike weights [wi(AIC)] (Burnham & Anderson, 2004), in order to simplify the model and avoid overfitting (Hawkins, 2004).

## RESULTS

### Content of media reports

We collected and analysed 314 media reports published between 2010 and 2020, discussing 344 spider-related events attributable to 97 unique events (Figure 1b). The average (±s.d.) number of media reports discussing each event was 3.52 ± 6.72 (range 1–33). The two most discussed events were i) the story of a traffic warden from Terni who was supposedly bitten by a Mediterranean recluse spider in 2018, covered by 33 media reports; and ii) the story of a woman supposedly bitten in 2019 by a Mediterranean recluse spider while sunbathing in Collecchio, covered by 31 media reports. All other events were covered by 20 media reports or fewer.

Most media reports focused on *Loxosceles rufescens* (n= 230; 66.9%) and *Latrodectus tredecimguttatus* (n= 97; 27.3%). Other species – *Cheiracanthium punctorium* (n= 14), *Steatoda* sp. (n= 4), and unidentified (n= 2) – were poorly represented (5.8%) and so we merged these under the category “Others”. Reports on *L. tredecimguttatus* mostly discussed human-spider encounters (Figure 2a), e.g., a farmer spotting a black widow while working in his field or a tourist photographing the species during a hike. Conversely, reports on *L. rufescens* mostly referred to bites (real or otherwise), including three unverified fatal cases (see discussion). Most media reports contained one or more photographs of the species (n= 298; 86.6%; Figure 2c), whereas only ca. 10% of media reports contained photographs of the bite (n= 33) (Figure 2d). Expert were sporadically mentioned in media reports (Figure 2e) and sensationalistic contents were more frequent in media reports referring to *L. rufescens* rather than other species (Figure 2f).

### Quality of media reports

One or more error types were present in 73% of media reports (Figure 3). The distribution of errors varied, however, depending on the species: most media reports referring to *L. tredecimguttatus* and other species contained no errors, whereas most reports on *L. rufescens* contained one or more errors (Figure 3a). The most frequent errors referred to spider morphology and anatomy (55.3%), species photographs (28.4%), and systematics and taxonomy (25.8%). Errors referring to venom and other physiological aspects were present in 15% of media reports (Figure 3b–e).

**Figure 3.**
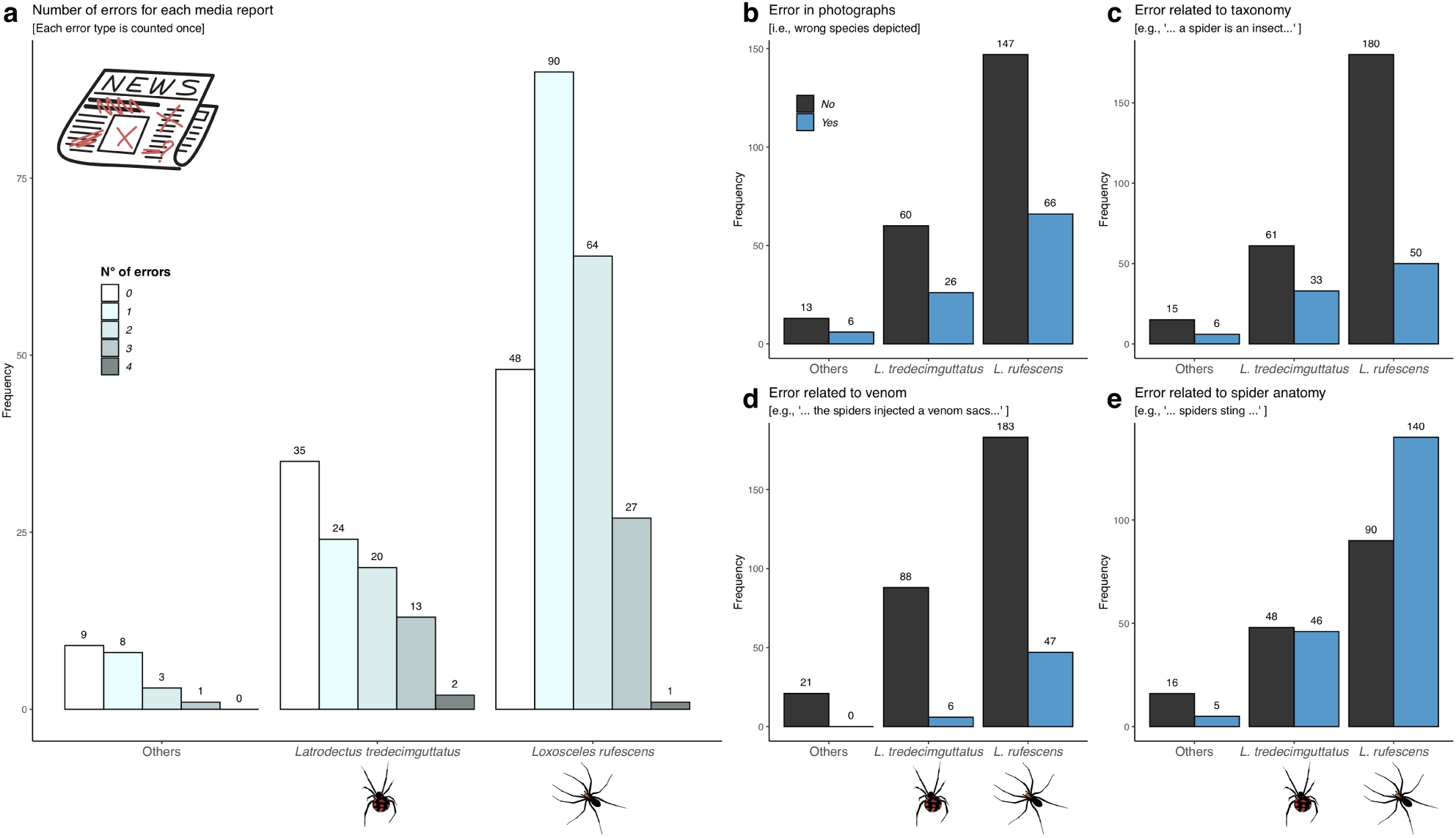
Quality of media reports: **a**) Total number of error for media reports focusing on *Latrodectus tredecimguttatus, Loxosceles rufescens*, and other species.

### Temporal distribution of media reports

We observed a strong temporal signal in the distribution of media reports between 2010 and 2019, with a recent increase in the number of news for both species, which was rather steadily increasing in *L. tredecimguttatus* and almost exponential in *L. rufescens* (Figure 4a). From a seasonal point of view (Figure 4b), we found that there was a clear summer peak, in July, in the frequency of reports for both species. This seasonal pattern was more evident for reports referring to *L. tredecimguttatus*.

**Figure 4.**
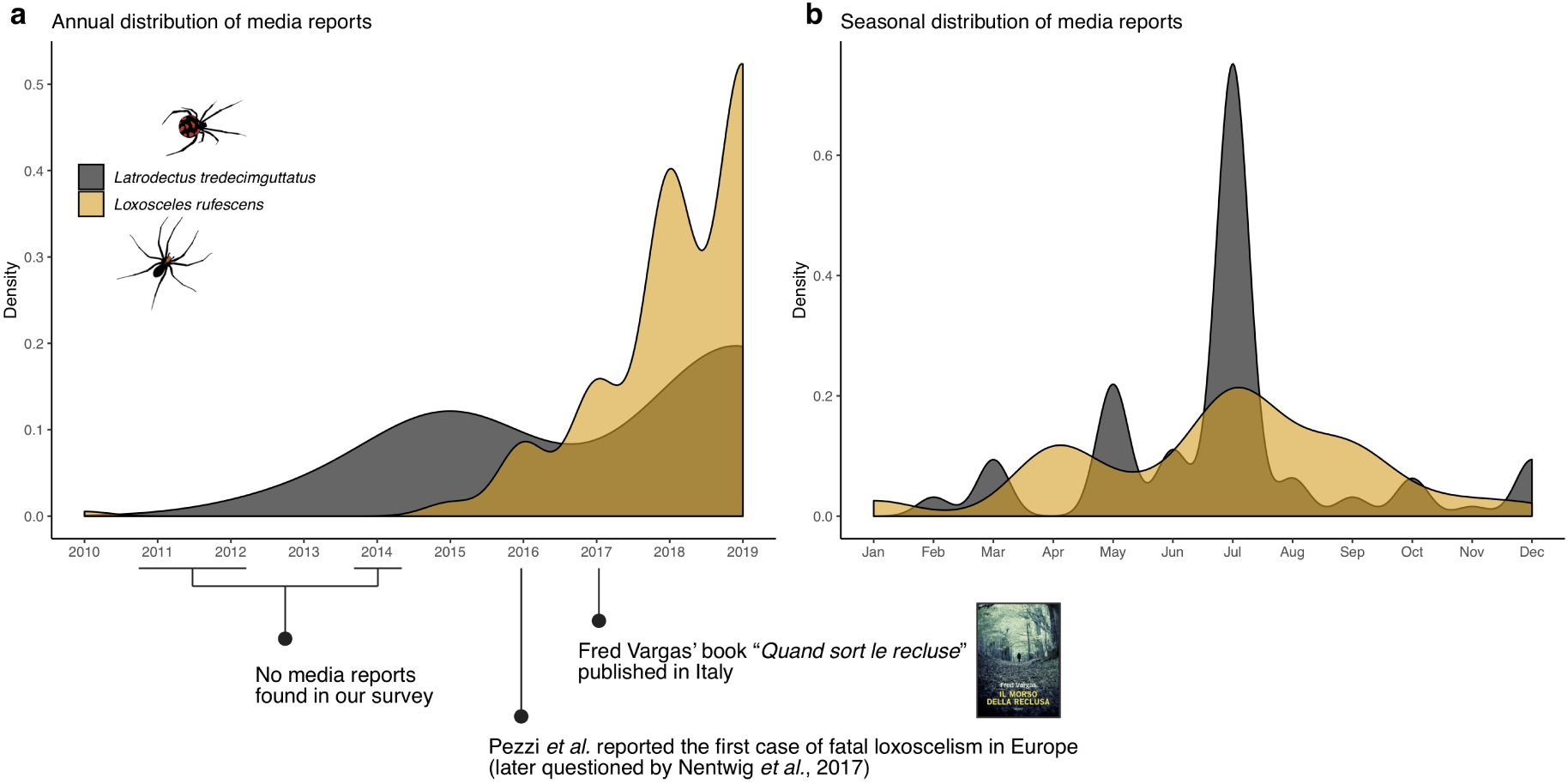
Temporal distribution of media reports: The cumulative curves for media reports referring to *Latrodectus tredecimguttatus* and *Loxosceles rufescens* are estimated with a kernel density. **a**) Annual distribution of media reports between 2010 and 2019. Few remarks are highlighted on the x-axis (see Discussion for details). **b**) Monthly distribution of media reports (cumulative of all years).

### Factors affecting the sharing of media reports on social media

The model that minimized AIC included year and sensationalism as fixed terms (Table 1). Random effect variance (± s.e.) was 6.55 ± 3.56 for Newspaper and 3.24 e^−5^ ± 0.01 for Event_ID. We found a significant positive effect of the year of publication, with recent media reports being, on average, more frequently shared on social media (Figure 5a). Furthermore, media reports with sensationalistic content were, on average, more frequently shared on social media (Figure 5b). All other factors had no significant influence on sharing on social media, and were discarded during model selection (Table 1).

**Table 1.** Result of model selection and estimated regression parameters. Estimated regression parameters (Estimated β ± S.E.) for fixed terms are given only for the selected model.

| Competing models | Estimated $\beta \pm \text{S.E.}$ | p | df | AIC | $\Delta\text{AIC}$ | $w_i$ |
| --- | --- | --- | --- | --- | --- | --- |
| Intercept | $-2688.36 \pm 434.44$ | - | 6 | 3057.76 | 0.0 | 0.414 |
| Year | $1.33 \pm 0.22$ | $5.6 \times 10^{-10}$ | | | | |
| Sensationalism | $1.15 \pm 0.50$ | 0.02 | | | | |
| Circulation + Year + Sensationalism | - | - | 7 | 3059.16 | 1.40 | 0.206 |
| Circulation + Year + Sensationalism + Expert opinion | - | - | 8 | 3059.72 | 1.96 | 0.156 |
| Circulation + Year + Sensationalism + Figure (bite) + Expert opinion | - | - | 9 | 3060.40 | 2.64 | 0.111 |
| Circulation + Year + Sensationalism + Figure (species) + Figure (bite) + Expert opinion | - | - | 10 | 3061.48 | 3.72 | 0.065 |
| Circulation + Year + Sensationalism + Species + Figure (species) + Figure (bite) + Expert opinion | - | - | 11 | 3063.16 | 5.40 | 0.028 |
| Event type + Circulation + Year + Sensationalism + Species + Figure (species) + Figure (bite) + Expert opinion | - | - | 12 | 3064.68 | 6.92 | 0.013 |
| Event type + Circulation + Year + Month + Month<sup>2</sup> + Sensationalism + Species + Figure (species) + Figure (bite) + Expert opinion | - | - | 14 | 3065.66 | 7.90 | 0.008 |

AIC= Akaike Information Criterion; $\Delta\text{AIC}$ = (AIC of the model – AIC of the best model); df= Degrees of freedom; p= *p*-value; $w_i$ = Akaike weights.

**Figure 5.**
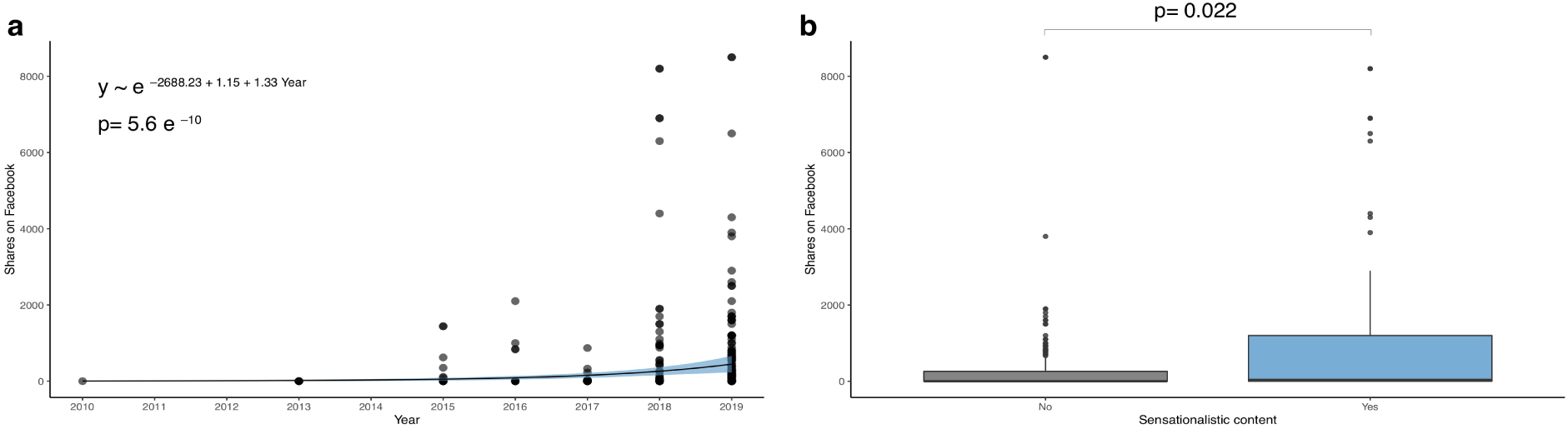
Factors driving the spreading of media reports on social media: The results are based on the most appropriate generalised linear mixed model (see Table 1 for model selection and estimated regression parameters). **a)** Predicted relationship between the number of Facebook shares and the year of publication of the media report. To generate the prediction, the effect of all factorial terms was summed to the intercept. **b**) Boxplots showing the difference between number of Facebook shares in neutral versus sensationalistic media reports.

## DISCUSSION

### Content of media reports and temporal distribution

We found that the quality of online newspaper articles focusing on spiders in Italy is, in general, rather poor. Media report quality appears to be independent of the newspaper’s circulation (national versus regional). Over 70% of media reports contained errors, 32% were characterized by an overstated content, and in virtually none of them was an expert consulted or interviewed. The two most represented species in the media reports were *Latrodectus tredecimguttatus* and *L. rufescens*, two species widely distributed in Italy (Pantini & Isaia, 2019). These two species belong to the only globally distributed genera responsible for medically important clinical syndromes, namely latrodectism and loxoscelism (Isbister & Fan, 2011). The fact that these species can deliver potentially harmful bites to humans seemingly explains why they are able to attract this great attention from the media.

*Loxosceles rufescens* is native to the Mediterranean basin (Planas, Saupe, Lima-Ribeiro, Peterson, & Ribera, 2014), but has been introduced to many areas of the world where it is considered an important invasive species (Nentwig, Pantini, & Vetter, 2017; Taucare-Rios, Nentwig, Bizama, & Bustamante, 2018). The Mediterranean recluse is a rather common inhabitant of natural and in-door habitats in Italy and thus, it seems likely that it has been coexisting with humans for centuries. Indeed, the species has been known in Italy since the second half of the XIX century, when the first catalogue on Italian spiders was published (Canestrini & Pavesi, 1868). According to scientific literature on Italian spiders (Pantini & Isaia, 2019), records of *L. rufescens* in indoor habitats have been increasing since 2000, with only one record before 1900, four between 1960 and 2000, and seven after 2000. Yet, the species began appearing in the media spotlight only in the past five years (Figure 4a). The increase of reports, often of poor quality and with a highly sensationalistic content, started just after the publication of the first supposed case of fatal loxoscelism in Europe (Pezzi et al., 2016; see discussion later). Coincidentally, this increase also came after the publication in Italy of *Quand sort la recluse*, a crime novel by Fred Vargas, where Chief inspector Jean-Baptiste Adamsberg has to deal with a series of murders committed using the venom of *L. rufescens*. While there is probably no causal relationship between these events and the increase in number of reports, it is interesting to note that several recent media reports in our database referenced both sources.

The quality of media reports referring to *L. tredecimguttatus* was better, and fewer reports had a sensationalistic content. *Latrodectus tredecimgutattus* was described based on specimens collected in Volterra (Tuscany). The species is distributed across a wide area in the Palearctic region, from the Mediterranean basin to Ukraine, Caucasus, Central Asia, and China (World Spider Catalog, 2020). In Italy, as well as in most in most other countries, the Mediterranean black widow is preferably found in ruderal areas of agricultural land and, just like *L. rufescens*, has been living close to humans for centuries. However, according to the data presented in scientific literature, its presence in strictly indoor habitats is infrequent – only 2 records out of the 23 available describing habitats (Pantini & Isaia, 2019). According to scientific literature on Italian spiders (Pantini & Isaia, 2019), most records of this species refer to natural or semi-natural (agricultural) habitats and only in one case (Pepe, 2005) has the species been reported in synanthropic habitats. The Mediterranean black widow began appearing in the media spotlight only in the past ten years (Figure 4a), with the highest number of media reports from late spring to early autumn, paralleling the period of highest activity of the species (Nentwig et al., 2020) and corresponding to the higher possibility of human-spider encounters. Given that most media reports on *L. tredecimguttatus* were in fact ‘Encounters’ (Figure 2a), namely reports of the species’ presence as provided by readers of the different newspapers, the distribution of news may be somehow tracking the species’ phenology, making it an unusual example of iEcology (Jarić et al., 2020).

We found that the risk scenario depicted by the media reports was unnecessarily alarmist, especially with regard to *L. rufescens*. First, no proven fatality due to a bite by *L. rufescens* has occurred globally (Nentwig et al., 2017). Second, overdiagnosis of spider bites is a rather common phenomenon for “popular” taxa such as *Loxosceles* and *Latrodectus* (Stuber & Nentwig, 2016). A conservative estimate would suggest that less than 10% of the bites reported in the media reports analysed here were delivered by the species described in the report (see Suchard, 2011). Third, in virtually none of the media reports is it written that the biting spider was brought to a hospital for identification, thus the causal attribution remains unconfirmed and merely suspected (Vetter & Isbister, 2008). Accordingly, the content of the majority of media reports analysed here has to be taken at best as anecdotic.

These considerations apply also to the three casualties associated with a bite by *L. rufescens* reported in the media reports. The only scientifically supported fatality refers to a case of loxoscelism in a woman, 65, dating back to 2015. This event was discussed in the medical literature (Pezzi et al., 2016), and later began to be mentioned by some journalists (n=7 media reports). However, the validity of this medical report was readily questioned by Nentwig et al. (2017), because the identity of the spider biting the woman was not ascertained. Allegedly: “*[The woman] was bitten the evening before hospitalization while cleaning the home cellar by a spider, which, from the description and place where the bite occurred, could probably be identified as the* Loxosceles rufescens *species*” (Pezzi et al., 2016). Two other fatalities covered in the media reports – Cagliari (2017) and Aosta (2020) – are unverifiable, and most likely wrong, given that neither was the bite ascertained nor was the spider collected and identified. The validity of these reports was even questioned in some newspapers, for example Report_ID 229 stating that “*The story of the men who died due to a violin spider bite is probably fake news*”, or Report_ID 115 observing that “*… he died three months after being stung* [bitten] *by a violin spider. But the cause of his death could be another*” (titles literally translated).

The seasonal pattern in the distribution of news with a marked summer peak (Figure 4b), parallels what was found by Cushing & Markwell (2010) when analysing newspapers articles on the Australian endemic Sydney funnel-web spider (*Atrax robustus*). The higher prevalence of secondary news during the summer holidays is a well known trend in journalism whereby, in the absence of more relevant news, a secondary subject such as a spider bite is frequently able to make it to the front pages.

### Social media amplification of sensationalistic contents

Social media have profoundly shaped the way information is produced and circulated. In recent years, social media platforms have become an important battlefield for political debates (Hall, Tinati, & Jennings, 2018), as well as the primary digital environments where people inform themselves and frame their perception of the world (Weeks, Ardèvol-Abreu, & Gil de Zúñiga, 2015). In parallel, social media have also become the preferential channel though which traditional news are disseminated and discussed (Lee & Ma, 2012), with most newspapers now actively using social media platforms such as Facebook and Twitter to spread their online contents more effectively (Ju, Jeong, & Chyi, 2014)

In line with this, we found that the share of spider-related news on social media has increased significantly in recent years (Figure 5a). In the contemporary era, where “*more iPhones are sold in a few days than there are tigers, elephants, and gorillas on the planet*” (Chapron, Levrel, Meinard, & Courchamp, 2018: p. 651), this result did not come as a surprise. However, not all news on spiders were shared with the same frequency online. While sensationalistic reports represent only about one third of the total media reports analysed in this survey, these were on average shared on Facebook two to three times more than neutral news (Figure 5b). This results is in accordance with general studies demonstrating that newspaper articles with content evoking strong positive or negative emotions are more likely to become viral (Berger & Milkman, 2012). Being shared on social media, sensationalistic news will inevitably be more widely read. Due to their sensationalistic content, they are also more likely to remain imprinted in a reader’s memory, especially in an arachnophobic reader’s, since it has been demonstrated that arachnophobics recall spider-relevant information more effectively (Smith-Janik & Teachman, 2008). On top of this, social media platforms are a fertile ground for emotional contagion, the phenomenon whereby emotional states are rapidly transferred to others leading to massive-scale emotional homogenisation (Kramer et al., 2014). This may contribute to empowering a biased perception of risk (Gerber et al., 2011) and facilitate the persistence of arachnophobic sentiments.

### Significance of results for spider conservation

Fear of spiders is one of the most prevalent animal-related phobias in humans (Mammola et al., 2017) and thus, spider-related contents are an effective emotional trigger (Smith-Janik & Teachman, 2008). We have shown how some journalists are able to exploit arachnophobic sentiments to their advantage, framing sensationalistic news capable of attracting substantial online attention. Sensationalistic news that dramatize and overstate the frequency of spiders “attacks” on humans are also those which most attract social media. Through emotional contagion, this biased representation is spread online. Ultimately, this may result in lowering public tolerance for spiders and lead to lower willingness for conservation and management efforts.

As demonstrated by Knight (2008), aesthetic and positive/negative features of animals correlate to the protection each taxon receives. Accordingly, the main challenges facing invertebrate conservationists is to change the perceived negative connotations of invertebrates by the public (Samways et al., 2020), raising awareness about the importance of these often uncharismatic organisms for the correct functioning of ecosystems (Cardoso et al., 2020). Spiders are apical predators in the invertebrate food web (Nyffeler & Birkhofer, 2017), while also representing a fundamental source of food for other organisms, such as birds. The importance of spiders has been even valued in economic terms, given that many species act as major biocontrol for pests in agroecosystem (Cotes et al., 2018; Michalko, Pekar, & Entling, 2019; Michalko, Pekár, Dul’a, & Entling, 2019), and their body structures, silk, and venom are constant sources for bio-inspired materials and engineering solutions (Hinman, Jones, & Lewis, 2000; Heim, Keerl, & Scheibel, 2009; Kang et al., 2014), as well as pharmaceutical products (Saez et al., 2010; Moore, Leung, Norton, & Cochran, 2013). Nevertheless, spiders are still largely underrepresented in global and regional conservation policies, particularly when compared to vertebrates (Leather, 2013; Davies et al., 2018; Fukushima, Mammola, & Cardoso, 2020) or charismatic insects such as butterflies and dragonflies (Milano et al., in prep.). In Europe, for example, spiders are almost entirely absent from international and national conservation policies, as well as from Italian legislation (Milano, Pantini, Mammola, & Isaia, 2017).

Traditional media have the potential to play an important role in changing the *status quo*, by offering the public unbiased representations of spiders. Thus, we urge journalists to renew their efforts toward objectivity and accuracy, which are best achieved by i) consulting and interviewing experts; ii) referring to scientific literature, as well as to modern online resources led by expert arachnologists (see, e.g., the “*Recluse or Not?*” project on Twitter; @RecluseOrNot); and iii) avoiding unmotivated sensationalism when describing biting events.

The traditional media arguably remain among the most powerful communication tools, capable of delivering their message effectively especially thanks to the aid of social media (Ju et al., 2014). If this potential is harnessed to the goal of delivering accurate information to the public at large (Papworth et al., 2015), this would facilitate the much-needed transition toward an unbiased protection of the diversity of life.

## Acknowledgements

Diego Fontaneto provided valuable suggestions. Special thanks are due to Stephen Cooper for proof-reading the English.

## Author contribution statement

Conceptualization: SM;

Data management: SM, VN, PP, and MI;

Data analysis: SM;

Writing, first draft: SM;

Writing, revisions: VN, PP, MI.

## Data availability statement

Data supporting this study will be deposited in a public online repository upon acceptance.

## LITERATURE CITED

Afshari, R. (2016). Bite like a spider, sting like a scorpion. Nature, 537(167). doi:10.1038/537167e

Bartoń, K. (2019). MuMIn: Multi-Model Inference.

Bennett, R. G., & Vetter, R. S. (2004). An approach to spider bites. Erroneous attribution of dermonecrotic lesions to brown recluse or hobo spider bites in Canada. Canadian Family Physician, 50, 1098–1101.

Berger, J., & Milkman, K. L. (2012). What Makes Online Content Viral? Journal of Marketing Research, 49(2), 192–205. doi:10.1509/jmr.10.0353

Bertone, M. A., Leong, M., Bayless, K. M., Malow, T. L. F., Dunn, R. R., & Trautwein, M. D. (2016). Arthropods of the great indoors: characterizing diversity inside urban and suburban homes. PeerJ, 4, e1582. doi:10.7717/peerj.1582

Blasco-Moreno, A., Pérez-Casany, M., Puig, P., Morante, M., & Castells, E. (2019). What does a zero mean? Understanding false, random and structural zeros in ecology. Methods in Ecology and Evolution, 10(7), 949–959. doi:10.1111/2041-210X.13185

Bombieri, G., Nanni, V., Delgado, M. del M., Fedriani, J. M., López-Bao, J. V., Pedrini, P., & Penteriani, V. (2018). Content Analysis of Media Reports on Predator Attacks on Humans: Toward an Understanding of Human Risk Perception and Predator Acceptance. BioScience, 68(8), 577–584. doi:10.1093/biosci/biy072

Burnham, K. P., & Anderson, D. R. (2004). *Multimodel inference: A Practical Information-Theoretic Approach*. (Springer-Verlag, Ed.), Sociological Methods and Research. New York. doi:10.1177/0049124104268644

Canestrini, G., & Pavesi, P. (1868). Araneidi italiani. Atti Della Società Italiana Di Scienze Naturali, Milano, 11(3), 738–872.

Cardoso, P., Barton, P. S., Birkhofer, K., Chichorro, F., Deacon, C., Fartmann, T., … Samways, M. J. (2020). Scientists’ warning to humanity on insect extinctions. Biological Conservation, 242, 108426. doi:10.1016/J.BIOCON.2020.108426

Cavell, M. (2018). Arachnophobia and early english literature. In L. Ashe, P. Knox, D. Lawton, & W. Scase (Eds.), New Medieval Literature (pp. 1–44). Cambridge: D.S. Brewer.

Chapron, G., Levrel, H., Meinard, Y., & Courchamp, F. (2018). A Final Warning to Planet Earth. Trends in Ecology & Evolution, 33(9), 651–652. doi:10.1016/j.tree.2017.12.010

Cotes, B., Gonzalez, M., Benitez, E., De Mas, E., Clemente-Orta, G., Campos, M., & Rodriguez, E. (2018). Spider Communities and Biological Control in Native Habitats Surrounding Greenhouses. INSECTS, 9(1). doi:10.3390/insects9010033

Cushing, N., & Markwell, K. (2010). ‘Watch out for these KILLERS!’: newspaper coverage of the Sydney funnel web spider and its impact on antivenom research. Health and History, 12(2), 79–96.

Davey, G. C. L. (1994). The ‘Disgusting’ Spider: The Role of Disease and Illness in the Perpetuation of Fear of Spiders. Society & Animals, 2(1), 17–25.

Davey, G. C. L., McDonald, A. S., Hirisave, U., Prabhu, G. G., Iwawaki, S., Jim, C. I., … C. Reimann, B. (1998). A cross-cultural study of animal fears. Behaviour Research and Therapy, 36 (7–8), 735–750. doi:10.1016/S0005-7967(98)00059-X

Davies, T., Cowley, A., Bennie, J., Leyshon, C., Inger, R., Carter, H., … Gaston, K. (2018). Popular interest in vertebrates does not reflect extinction risk and is associated with bias in conservation investment. PLOS ONE, 13(9), e0203694. Retrieved from https://doi.org/10.1371/journal.pone.0203694

DeLoache, J. S., Pickard, M. B., & LoBue, V. (2010). How very young children think about animals. In How animals affect us: Examining the influences of human–animal interaction on child development and human health. doi:10.1037/12301-004

Diaz, J. H., & Leblanc, K. E. (2007). Common spider bites. American Family Physician, 75(6), 869–873.

Drijfhout, M., Kendal, D., & Green, P. T. (2020). Understanding the human dimensions of managing overabundant charismatic wildlife in Australia. Biological Conservation, 244, 108506. doi:10.1016/J.BIOCON.2020.108506

Fournier, D. A., Skaug, H. J., Ancheta, J., Ianelli, J., Magnusson, A., Maunder, M. N., … Sibert, J. (2012). AD Model Builder: Using automatic differentiation for statistical inference of highly parameterized complex nonlinear models. Optimization Methods and Software, 27, 233–249. doi:10.1080/10556788.2011.597854

Frank, J., Johansson, M., & Flykt, A. (2015). Public attitude towards the implementation of management actions aimed at reducing human fear of brown bears and wolves. Wildlife Biology, 21, 122–130. doi:10.2981/wlb.13116

Franklin, A., & White, R. (2001). Animals and modernity: Changing human–animal relations, 1949–98. Journal of Sociology, 37(3), 219–238. doi:10.1177/144078301128756319

Fukushima, C. S., Mammola, S., & Cardoso, P. (2020). Global wildlife trade permeates the Tree of Life. Biological Conservation, in press. doi:10.1016/j.biocon.2020.108503

Gerber, D. L. J., Burton-Jeangros, C., & Dubied, A. (2011). Animals in the media: New boundaries of risk? Health, Risk and Society, 13(1), 17–30. doi:10.1080/13698575.2010.540646

Gerdes, A. B. M., Uhl, G., & Alpers, G. W. (2009). Spiders are special: fear and disgust evoked by pictures of arthropods. Evolution and Human Behavior, 30(1), 66–73. doi:10.1016/J.EVOLHUMBEHAV.2008.08.005

Hall, W., Tinati, R., & Jennings, W. (2018). From Brexit to Trump: Social media’s role in democracy. Computer, 51(1), 18–27. doi:10.1109/MC.2018.1151005

Hathaway, R. S., Bryant, A. E. M., Draheim, M. M., Vinod, P., Limaye, S., & Athreya, V. (2017). From fear to understanding: Changes in media representations of leopard incidences after media awareness workshops in mumbai, India. Journal of Urban Ecology, 3(1), jux009. doi:10.1093/jue/jux009

Hauke, T. J., & Herzig, V. (2017). Dangerous arachnids—Fake news or reality? Toxicon, 138, 173–183. doi:10.1016/J.TOXICON.2017.08.024

Hawkins, D. M. (2004). The Problem of Overfitting. Journal of Chemical Information and Computer Sciences, 44, 1–12. doi:10.1021/ci0342472

Heim, M., Keerl, D., & Scheibel, T. (2009). Spider silk: From soluble protein to extraordinary fiber. Angewandte Chemie - International Edition, 48(20), 3584–3596. doi:10.1002/anie.200803341

Hicks, J. R., & Stewart, W. P. (2018). Exploring potential components of wildlife-inspired awe. Human Dimensions of Wildlife, 23(3), 293–295. doi:10.1080/10871209.2018.1419518

Hinman, M. B., Jones, J. A., & Lewis, R. V. (2000). Synthetic spider silk: A modular fiber. Trends in Biotechnology, 18(9), 374–379. doi:10.1016/S0167-7799(00)01481-5

Isbister, G. K., & Fan, H. W. (2011). Spider bite. The Lancet, 378, 2039–2047. doi:10.1016/S0140-6736(10)62230-1

Jacobs, M. H. (2009). Why Do We Like or Dislike Animals? Human Dimensions of Wildlife, 14(1), 1–11. doi:10.1080/10871200802545765

Jacobs, M. H. (2012). Human Emotions Toward Wildlife. Human Dimensions of Wildlife, 17(1), 1–3. doi:10.1080/10871209.2012.653674

Jambrina, C. U., Vacas, J. M., & Sánchez-Barbudo, M. (2010). Preservice teachers’ conceptions about animals and particularly about spiders. Electronic Journal of Research in Educational Psychology, 8(2), 787–814.

Jarić, I., Correia, R. A., Brook, B. W., Buettel, J. C., Courchamp, F., Di Minin, E., … Roll, U. (2020). iEcology: Harnessing Large Online Resources to Generate Ecological Insights. Trends in Ecology & Evolution. doi: https://doi.org/10.1016/j.tree.2020.03.003

Jones, S. (2006). A political ecology of wildlife conservation in Africa. Review of African Political Economy, 33(109), 483–495. doi:10.1080/03056240601000911

Ju, A., Jeong, S. H., & Chyi, H. I. (2014). Will Social Media Save Newspapers: Examining the effectiveness of Facebook and Twitter as news platforms. Journalism Practice, 8, 1–17. doi:10.1080/17512786.2013.794022

Kang, D., Pikhitsa, P. V., Choi, Y. W., Lee, C., Shin, S. S., Piao, L., … Choi, M. (2014). Ultrasensitive mechanical crack-based sensor inspired by the spider sensory system. Nature, 516(7530), 222–226. doi:10.1038/nature14002

Knight, A. J. (2008). ‘Bats, snakes and spiders, Oh my!’ How aesthetic and negativistic attitudes, and other concepts predict support for species protection. Journal of Environmental Psychology, 28(1), 94–103. doi:10.1016/j.jenvp.2007.10.001

Knopff, A. A., Knopff, K. H., & St. Clair, C. C. (2016). Tolerance for cougars diminished by high perception of risk. Ecology and Society, 21, 33. doi:10.5751/ES-08933-210433

Kramer, A. D. I., Guillory, J. E., & Hancock, J. T. (2014). Experimental evidence of massive-scale emotional contagion through social networks. Proceedings of the National Academy of Sciences, 111(24), 8788 LP–8790. doi:10.1073/pnas.1320040111

Langley, R. L. (2005). Animal-Related Fatalities in the United States—An Update. Wilderness & Environmental Medicine, 16(2), 67–74. doi:https://doi.org/10.1580/1080-6032(2005)16[67:AFITUS]2.0.CO;2

Leather, S. R. (2013). Institutional vertebratism hampers insect conservation generally; not just saproxylic beetle conservation. Animal Conservation, 16, 379–380. doi:10.1111/acv.12068

Lee, C. S., & Ma, L. (2012). News sharing in social media: The effect of gratifications and prior experience. Computers in Human Behavior, 28, 331–339. doi:10.1016/j.chb.2011.10.002

Lemelin, R. H., & Yen, A. (2015). Human-Spider Entanglements: Understanding and Managing the Good, the Bad, and the Venomous. Anthrozoös, 28(2), 215–228. doi:10.1080/08927936.2015.11435398

Mammola, S., Michalik, P., Hebets, E. A., & Isaia, M. (2017). Record breaking achievements by spiders and the scientists who study them. PeerJ, 5(10), e3972. doi:10.7717/peerj.3972

Merckelbach, H., Muris, P., & Schouten, E. (1996). Pathways to fear in spider phobic children. Behaviour Research and Therapy, 34(11–12), 935–938. doi:10.1016/S0005-7967(96)00052-6

Michalko, R., Pekár, S., Dul’a, M., & Entling, M. H. (2019). Global patterns in the biocontrol efficacy of spiders: A meta-analysis. Global Ecology and Biogeography, 28(9), 1366–1378. doi:10.1111/geb.12927

Michalko, R., Pekar, S., & Entling, M. H. (2019). An updated perspective on spiders as generalist predators in biological control. OECOLOGIA, 189(1), 21–36. doi:10.1007/s00442-018-4313-1

Michalski, K., & Michalski, S. (2010). Spider. London, UK: Reaktion Books Ltd. Milano, F., et al. (in prep.) Spider conservation in Europe.

Milano, F., Pantini, P., Mammola, S., & Isaia, M. (2017). La conservazione dell’araneofauna in Italia e in Europa [Spider conservation in Italy and Europe]. Atti Accademia NazionaleItaliana Di Entomologia, 65, 91–103.

Moore, S. J., Leung, C. L., Norton, H. K., & Cochran, J. R. (2013). Engineering Agatoxin, a Cystine-Knot Peptide from Spider Venom, as a Molecular Probe for In Vivo Tumor Imaging. PLOS ONE, 8(4), e60498. Retrieved from https://doi.org/10.1371/journal.pone.0060498

Nanni, V., Caprio, E., Bombieri, G., Schiaparelli, S., Chiorri, C., Mammola, S., … Penteriani, V. (2020). Social Media and Large Carnivores: Sharing Biased News on Attacks on Humans. Frontiers in Ecology and Evolution, 8, 71. doi:10.3389/fevo.2020.00071

Nentwig, W., Blick, T., Bosmans, R., Gloor, D., Hänggi, A., & Kropf, C. (2020). Araneae - Spider of Europe. doi:https://doi.org/10.24436/1

Nentwig, W., Gnädinger, M., Fuchs, J., & Ceschi, A. (2013). A two year study of verified spider bites in Switzerland and a review of the European spider bite literature. Toxicon, 73, 104–110. doi:10.1016/j.toxicon.2013.07.010

Nentwig, W., & Kuhn-Nentwig, L. (2013). Spider venoms potentially lethal to humans. In W. Nentwig (Ed.), Spider Ecophysiology. Heidelberg: Springer. doi:10.1007/978-3-642-33989-9_19

Nentwig, W., Pantini, P., & Vetter, R. S. (2017). Distribution and medical aspects of Loxosceles rufescens, one of the most invasive spiders of the world (Araneae: Sicariidae). Toxicon, 132, 19–28. doi:10.1016/j.toxicon.2017.04.007

Nyffeler, M., & Birkhofer, K. (2017). An estimated 400–800 million tons of prey are annually killed by the global spider community. The Science of Nature, 104(3), 30. doi:10.1007/s00114-017-1440-1

Pantini, P., & Isaia, M. (2019). Araneae.it: the online Catalog of Italian spiders, with addenda on other Arachnid Orders occurring in Italy (Arachnida: Araneae, Opiliones, Palpigradi, Pseudoscorpionida, Scorpiones, Solifugae). Online at www.araneae.it, accessed on {26 April 2020}. Fragmenta Entomologica, 52(2), 127–152. doi:10.4081/fe.2019.374

Papworth, S. K., Nghiem, T. P. L., Chimalakonda, D., Posa, M. R. C., Wijedasa, L. S., Bickford, D., & Carrasco, L. R. (2015). Quantifying the role of online news in linking conservation research to Facebook and Twitter. Conservation Biology, 29(3), 825–833. doi:10.1111/cobi.12455

Pepe, R. (2005). Basi zoologiche-naturalistiche del tarantismo nel Salento. Thalassia Salentina, 27(2004), 47–62.

Pezzi, M., Giglio, A. M., Scozzafava, A., Filippelli, O., Serafino, G., & Verre, M. (2016). Spider Bite: A Rare Case of Acute Necrotic Arachnidism with Rapid and Fatal Evolution. Case Reports in Emergency Medicine, 2016, 7640789. doi:10.1155/2016/7640789

Planas, E., Saupe, E. E., Lima-Ribeiro, M. S., Peterson, A. T., & Ribera, C. (2014). Ecological niche and phylogeography elucidate complex biogeographic patterns in Loxosceles rufescens (Araneae, Sicariidae) in the Mediterranean Basin. BMC EVOLUTIONARY BIOLOGY, 14. doi:10.1186/s12862-014-0195-y

Prokop, P., & Tunnicliffe, S. D. (2008). ‘Disgusting’ animals: Primary school children’s attitudes and myths of bats and spiders. Eurasia Journal of Mathematics, Science and Technology Education, 4(2), 87–97. doi:10.12973/ejmste/75309

R Core Team. (2018). R: A Language and Environment for Statistical Computing. Vienna, Austria: R Foundation for Statistical Computing.

Saez, N. J., Senff, S., Jensen, J. E., Er, S. Y., Herzig, V., Rash, L. D., & King, G. F. (2010). Spider-venom peptides as therapeutics. Toxins, 2(12), 2851–2871. doi:10.3390/toxins2122851

Samways, M. J., Barton, P. S., Birkhofer, K., Chichorro, F., Deacon, C., Fartmann, T., … Cardoso, P. (2020). Solutions for humanity on how to conserve insects. Biological Conservation, 242, 108427. doi:10.1016/J.BIOCON.2020.108427

Savage, I. (2013). Comparing the fatality risks in United States transportation across modes and over time. Research in Transportation Economics, 43, 9–22. doi:10.1016/j.retrec.2012.12.011

Simaika, J. P., & Samways, M. J. (2018). Insect conservation psychology. Journal of Insect Conservation, 22(3), 635–642. doi:10.1007/s10841-018-0047-y

Singh, S. (2009). Governing Anti-conservation Sentiments: Forest Politics in Laos. Human Ecology, 37(6), 749. doi:10.1007/s10745-009-9276-8

Slovic, P., & Peters, E. (2006). Risk perception and affect. Current Directions in Psychological Science, 15, 322–325. doi:10.1111/j.1467-8721.2006.00461.x

Smith-Janik, S. B., & Teachman, B. A. (2008). Impact of priming on explicit memory in spider fear. Cognitive Therapy and Research, 32(2), 291–302. doi:10.1007/s10608-007-9122-5

Straka, T. M., Miller, K. K., & Jacobs, M. H. (2020). Understanding the acceptability of wolf management actions: roles of cognition and emotion. Human Dimensions of Wildlife, 25(1), 33–46. doi:10.1080/10871209.2019.1680774

Strommen, E. (1995). Lions and tigers and bears, oh my! children’s conceptions of forests and their inhabitants. Journal of Research in Science Teaching, 32(7), 683–698. doi:10.1002/tea.3660320704

Stuber, M., & Nentwig, W. (2016). How informative are case studies of spider bites in the medical literature? Toxicon, 114, 40–44. doi:10.1016/j.toxicon.2016.02.023

Suchard, J. R. (2011). ‘Spider bite’ lesions are usually diagnosed as skin and soft-tissue infections. Journal of Emergency Medicine, 41(5), 473–481. doi:10.1016/j.jemermed.2009.09.014

Taucare-Rios, A., Nentwig, W., Bizama, G., & Bustamante, R. O. (2018). Matching global and regional distribution models of the recluse spider Loxosceles rufescens: to what extent do these reflect niche conservatism? MEDICAL AND VETERINARY ENTOMOLOGY, 32(4), 490–496. doi:10.1111/mve.12311

Turnbull, A. L. (1973). Ecology of the True Spiders (Araneomorphae). Annual Review of Entomology, 18(1), 305–348. doi:10.1146/annurev.en.18.010173.001513

Vetter, R. S. (2004). Myths about spider envenomations and necrotic skin lesions. Lancet, 364, 484–485. doi:10.1016/S0140-6736(04)16824-4

Vetter, R. S., Hinkle, N. C., & Ames, L. M. (2009). Distribution of the Brown Recluse Spider (Araneae: Sicariidae) in Georgia with Comparison to Poison Center Reports of Envenomations. Journal of Medical Entomology, 46(1), 15–20. doi:10.1603/033.046.0103

Vetter, R. S., & Isbister, G. K. (2008). Medical Aspects of Spider Bites. Annual Review of Entomology, 53, 409–429. doi:10.1146/annurev.ento.53.103106.093503

Vetter, R. S., Warrell, D. A., Isbister, G. K., White, J., Currie, B. J., & Bush, S. P. (2005). Spider bites: Addressing mythology and poor evidence. The American Journal of Tropical Medicine and Hygiene, 72(4), 361–364. doi:10.4269/ajtmh.2005.72.361

Vosoughi, S., Roy, D., & Aral, S. (2018). The spread of true and false news online. Science, 359(6380), 1146 LP–1151. doi:10.1126/science.aap9559

Weeks, B. E., Ardèvol-Abreu, A., & Gil de Zúñiga, H. (2015). Online Influence? Social Media Use, Opinion Leadership, and Political Persuasion. International Journal of Public Opinion Research, 29(2), 214–239. doi:10.1093/ijpor/edv050

White, J. (2003). Debunking spider bite myths. Medical Journal of Australia, 179(4), 180–181. doi:10.5694/j.1326-5377.2003.tb05493.x

Wickham, H. (2016). ggplot2: Elegant Graphics for Data Analysis. New York: Springer-Verlag.

Wilson, R. E., Gosling, S. D., & Graham, L. T. (2012). A Review of Facebook Research in the Social Sciences. Perspectives on Psychological Science, 7(3), 203–220. doi:10.1177/1745691612442904

World Spider Catalog. (2020). World Spider Catalog. Version 20.5. doi:10.24436/2

Zainal Abidin, Z. A., & Jacobs, M. (2019). Relationships between valence towards wildlife and wildlife value orientations. Journal for Nature Conservation, 49, 63–68. doi:10.1016/J.JNC.2019.02.007

Zuur, A. F., & Ieno, E. N. (2016). A protocol for conducting and presenting results of regression-type analyses. Methods in Ecology and Evolution, 7(6), 636–645. doi:10.1111/2041-210X.12577

